# Secure Attachment despite Childhood Maltreatment: Behavioral and Neural Correlates of Interpersonal Resilience

**DOI:** 10.64898/2026.07.20.739488

**Authors:** Katharina Brosch, Vincent Hammes, Paula Usemann, Frederike Stein, Sunny X. Tang, Rebecca M. Schwartz, Florian Thomas-Odenthal, Lea Teutenberg, Nooshin Javaheripour, Susanne Meinert, Dominik Grotegerd, Elisabeth J. Leehr, Tiana Borgers, Kira Flinkenflügel, Janik Goltermann, Anne Hildebrand, Sabrina Teckentrup, Roman Vogler, Tim Hahn, Udo Dannlowski, Philipp Kanske, Benjamin Straube, Hamidreza Jamalabadi, Andreas Jansen, Igor Nenadić, Tilo Kircher, Nina Alexander

## Abstract

**Background:** Childhood maltreatment is a traumatic interpersonal stressor that increases vulnerability for depression in adulthood. However, some individuals show secure attachment despite childhood maltreatment, a pattern that can be described as interpersonal resilience. The present study examined the behavioral and neural correlates of interpersonal resilience, defined as secure attachment in adults exposed to childhood maltreatment.

**Methods:** We analyzed structural 3T MRI data from 1,317 adults, including healthy participants and individuals with partially or fully remitted major depression. Gray matter volume was estimated from structural MRI data using voxel-based morphometry. Attachment style and childhood maltreatment were assessed using the Relationship Scales Questionnaire and the Childhood Trauma Questionnaire, respectively. A 2×2 design (childhood maltreatment by attachment style) tested main and interaction effects on behavioral outcomes and brain structure.

**Results:** Interpersonally resilient individuals with secure attachment and maltreatment reported significantly better mental health outcomes compared to insecurely attached adults with maltreatment. Differences included lower self-reported and rater-based depressive symptoms, lower global symptom severity, and higher global functioning. In the neuroimaging analyses, we identified a significant childhood maltreatment by attachment style interaction in the left supramarginal gyrus, with larger gray matter volume in resilient individuals compared to all other groups. This effect remained robust across multiple sensitivity analyses, controlling for medication load, antidepressant intake, diagnosis group, as well as in a complementary dimensional analysis.

**Conclusions:** The results identify a potential neural correlate of interpersonal resilience. Larger gray matter volume in the left supramarginal gyrus, a region previously implicated in perspective taking and self-other distinction among other functions, may be relevant to more adaptive interpersonal functioning after early adversity. Together with the robust behavioral effects, these findings are consistent with secure attachment as a protective factor that may be associated with attenuated effects of childhood maltreatment on mental health.

## Main

Attachment theory provides a framework for understanding how individuals navigate close relationships and regulate stress. Secure attachment is characterized by positive internal models of self and others, which enable interdependence and foster adaptive emotion regulation, effective help-seeking, and the ability to maintain stable, supportive relationships ^1,2^. These skills, in turn, are protective against psychopathology and are consistently linked to higher well-being, life satisfaction, and fewer depressive symptoms ^3–5^.

Insecure attachment styles, which include anxious-preoccupied, dismissive-avoidant, and disorganized attachment styles, reflect disruptions in the internal models of self and others and have been identified as transdiagnostic vulnerability factors across psychiatric disorders ^6^. In relationships, these insecure styles can manifest as excessive reassurance- seeking and fear of abandonment (anxious-preoccupied), discomfort with closeness and compulsive self-reliance (dismissive-avoidant), or contradictory approach-avoidance behaviors toward attachment figures (disorganized) ^1^.

One major developmental context shaping attachment security is exposure to early interpersonal adversity. Childhood maltreatment, which includes experiences of abuse and neglect, is among the strongest known risk factors for the development of major depressive disorder (MDD) and for unfavorable illness trajectories including earlier onset, chronic course, and poorer treatment response ^7–9^. As a form of interpersonal trauma, childhood maltreatment can disrupt fundamental expectations of safety, trust, and relational security ^10^, thereby increasing the likelihood of attachment insecurity later in life. Consistent with this, childhood maltreatment is negatively associated with secure attachment and positively associated with insecure attachment styles ^11,12^. Attachment insecurity has in turn been proposed as a key psychological mechanism linking early adversity to later psychopathology. Accordingly, several studies have demonstrated indirect pathways from childhood maltreatment to depressive and psychotic symptoms through attachment insecurity ^13–17^.

However, attachment insecurity is not an inevitable outcome of childhood maltreatment. Some individuals exposed to early adversity nevertheless maintain secure attachment, potentially reflecting a form of interpersonal resilience. Here, we use the term interpersonal resilience to describe this pattern of adaptive interpersonal functioning despite a history of maltreatment, operationalized as low fear of abandonment and low fear of closeness ^3^. Understanding this protective process is crucial for explaining why individuals with similar adversity histories show different psychological outcomes ^18^.

At the neural level, both childhood maltreatment and attachment style have been associated with alterations in brain structure and function. Childhood maltreatment has been linked to widespread alterations in regions supporting emotion regulation, self-referential processing, and social cognition, including the prefrontal cortex, amygdala, hippocampus, and parietal areas ^19,20^. In parallel, studies in healthy samples suggest associations between attachment dimensions and gray matter volume (GMV), including regions such as the insula, pars opercularis, and anterior cingulate gyrus, related to social and emotional processing ^21–23^.

However, little is known about how attachment security and childhood maltreatment jointly relate to brain structure. It remains unclear whether secure attachment in individuals with a history of childhood maltreatment, conceptualized here as interpersonal resilience, is associated with specific neuroanatomical alterations.

Building on the concept of interpersonal resilience, we examined whether secure attachment buffers the association between childhood maltreatment and depressive symptoms and whether such resilience-related patterns are also reflected in brain structure. To this end, we analyzed data from a large sample of 1,317 adults. In line with a dimensional perspective on depression, the sample included individuals with and without a history of depressive disorder, allowing us to capture variability in depressive symptoms across the population rather than focusing solely on categorical patient-control comparisons. This approach is especially relevant because attachment insecurity and depressive symptoms are not confined to clinical thresholds but vary continuously across the population, and even subthreshold depressive symptom levels have been linked to alterations in brain structure^24,25^.

Participants with acute depressive episodes were excluded to minimize potential state-related confounds in the assessment of attachment style. Using voxel-based morphometry of high-resolution structural MRI, we tested whether attachment security moderates the association between childhood maltreatment and gray matter volume across the brain. Behaviorally, we hypothesized that individuals with a history of childhood maltreatment who maintain secure attachment would exhibit lower depressive symptom severity and higher global functioning than maltreated individuals with insecure attachment. On the neural level, we expected interpersonal resilience to be associated with larger gray matter volume in regions supporting social cognition and emotion regulation. As attachment security can be strengthened through therapeutic and relational processes, identifying neural correlates of this kind of resilience may help pinpoint who benefits most from attachment- focused interventions and provide objective markers to track their effects ^26–28^.

## Methods and Materials

### Participants

We included *N* = 1,317 participants aged 18-65 years, comprising individuals with fully or partially remitted MDD and psychiatrically healthy individuals. Participants were part of the bicentric German Marburg-Münster Affective Cohort Study (MACS), part of the FOR2107 consortium investigating the neurobiology of affective and psychotic disorders ^29^.

General exclusion criteria of the MACS cohort study included history of neurological conditions, severe medical conditions (e.g., cancer, multiple sclerosis), current or lifetime alcohol dependency, current substance dependency, current use of benzodiazepines, and IQ ≤ 80 estimated using the multiple-choice vocabulary test ^30^. In the MACS cohort study, any current or past psychiatric disorder, or lifetime intake of psychotropic medication constituted exclusion criteria for the healthy control (HC) group.

The study protocols were approved by the local Ethics Committees of Marburg and Münster, Germany, according to the Declaration of Helsinki. Written informed consent was obtained from all participants before participation. Participants were recruited from the area of Marburg and Münster, Germany, from in and out-patient departments of the respective universities, local psychiatric hospitals, psychotherapists’ offices, and through advertisements. Participation in the study was financially compensated and included MRI scanning, a battery of neuropsychological testing, self-report questionnaires, rater-based scales, biomaterial collection, and a semi-structured clinical interview for DSM-IV-TR (SCID-I) administered by trained raters ^31^.

In the current study, we included participants with and without a history of MDD, assessed using SCID-I. We excluded participants in an acute major depressive episode, and participants with rater-based Hamilton depression rating (HAMD-17) scores > 16 (indicating moderate/severe MDD) ^32,33^.

### Assessment of childhood maltreatment

Childhood maltreatment was assessed using the 28-item self-report Childhood Trauma Questionnaire (CTQ), consisting of five subscales of maltreatment (physical, emotional and sexual abuse, and physical and emotional neglect), with five items per scale, as well as three minimization and denial items ^34^. The German version has demonstrated good psychometric properties ^35^. Individuals were classified as maltreated when they met at least one of the validated cut-off scores of the five subscales ^36^.

### Assessment of adult attachment style

Attachment style was assessed using the 30-item self-report Relationship Scales Questionnaire (RSQ) ^37^. The German version was validated by Steffanowski et al., (2001) and identifies four factors: fear of closeness (avoidance), fear of abandonment (anxiety), lack of trust, and independence. Following Steffanowski et al., the first two factors (fear of closeness and fear of abandonment) can be used to categorize individuals into secure and insecure attachment styles (insecure fearful, insecure preoccupied, insecure dismissing). For the present study, we calculated sum scores for the avoidance and anxiety scales and applied the corresponding cut-off scores to classify participants dichotomously as securely or insecurely attached.

### Behavioral analyses

To investigate whether interpersonally resilient individuals (i.e., securely attached despite childhood maltreatment) differed in behavioral domains relevant to mental health, we examined self-rated depressive symptoms using the Beck Depression Inventory (BDI) ^39^ and clinician-rated depressive symptoms using the Hamilton Depression Rating Scale (HAM-D)^33^. Additional measures included perceived stress (Perceived Stress Scale; PSS) ^40^, self- reported resilience (Resilience Scale; RS-25) ^41^, global symptom severity (Symptom Checklist-90; SCL-90-R) ^42^, and global functioning (Global Assessment of Functioning; GAF)^43^.

### MRI data acquisition and preprocessing

To assess effects of interpersonal resilience on GMV, MRI scans (T1-weighted images) were acquired using 3T scanners in Marburg and Münster, Germany, using a fast gradient echo MP-RAGE sequence (voxel size of 1 × 1 × 1 mm). The following parameters were applied: Marburg: Tim Trio scanner, Siemens; 12-channel head matrix Rx-coil, 176 sagittal slices, repetition time (TR) = 1.9 s, echo time(TE) = 2.26 ms, inversion time (TI) = 900 ms, flip angle = 9°; Münster: Prisma scanner, Siemens, 20- channel head matrix Rx-coil, 192 sagittal slices, TR = 2.13 s, TE = 2.28 ms, TI = 900 ms, flip angle = 8°. Data acquisition and quality control followed the procedures described in Vogelbacher et al., (2018).

All scans were visually inspected by a senior clinician (UD) and scans with strong artifacts or anatomical abnormalities were excluded before preprocessing. Structural MRI data were processed using a voxel-based morphometry (VBM) approach as implemented in the CAT12-Toolbox (Computational Anatomy Toolbox for SPM, build 1720, Structural Brain Mapping group, Jena University Hospital, Germany) (http://dbm.neuro.uni-jena.de/cat/) building on SPM12 (Statistical Parametric Mapping, Institute of Neurology, London, UK) using default parameters. Preprocessing steps included image segmentation (gray matter, white matter, cerebrospinal fluid), spatial registration, and normalization. Data were normalized to Montreal Neurological Institute (MNI) space. During preprocessing, total intracranial volume (TIV) was calculated. Data were smoothed using a Gaussian kernel of 8mm full width at half maximum.

### Statistical analyses

For phenotypical and brain structural analyses, we applied a 2 x 2 design (childhood maltreatment x attachment style) and investigated main and interaction effects. Participants were categorized into four groups based on attachment security (secure vs. insecure) and childhood maltreatment (present vs. absent). We included age, sex, site and, in the case of brain structural analyses, TIV and body coil change as covariates, in line with our quality assurance protocol ^44^.

For behavioral analyses, SPSS 27 (Statistical Package for the Social Sciences, IBM) and R (version 4.5.1) within the R Studio environment (version 2026.1.0.392) were used.

As behavioral measures were moderately correlated (see Supplementary Figure 1), we applied MANCOVA to jointly investigate effects of childhood maltreatment x attachment on BDI-I, HAM-D, PSS, RS-25, GAF, and SCL-90-R global symptom severity index.

For the brain structural analyses, a whole-brain 2 x 2 model was set up in SPM12 (building on SPM12, Statistical Parametric Mapping, Wellcome Trust Centre for Neuroimaging, London, UK, running under Matlab, R2017a). As recommended by CAT12, we applied absolute threshold masking with a value of 0.1. Results were considered significant at *p* < 0.05 cluster-level with family wise error-correction (FWE). Significant clusters were labelled using the Dartel Neuromorphometrics atlas (http://www.neuromorphometrics.com/).

To test the robustness of the observed interaction effect, additional sensitivity analyses were conducted using the extracted subject-level eigenvariate values of the cluster. In these follow-up models, antidepressant intake (dichotomous) and medication load (Sackeim Index) were separately included as additional covariates in the MDD subgroup. Further, diagnostic group (healthy controls vs. MDD) was included as an additional covariate in the full sample. Finally, to complement the categorical CM × attachment analysis and examine whether the observed effects held across the full range of attachment security, rather than only between discrete groups, we performed a dimensional moderation analysis using the extracted eigenvariate values from the significant GMV cluster. Using linear regression, we tested how continuous childhood maltreatment severity (CTQ sum score) interacted with continuous attachment insecurity (multiplication term of RSQ subscales fear of closeness and fear of abandonment) to predict cluster eigenvariate values while controlling for age, sex, site, and TIV.

## Results

### Sample characteristics

Descriptive statistics for the four study groups are presented in Table 1 (InsecCM− n = 270; InsecCM+ n = 300; SecCM− n = 494; SecCM+ n = 253). The groups differed significantly in age (p < .001) and diagnostic status (p < .001), whereas sex distribution did not differ across groups (p = .300). As expected, individuals classified as maltreated showed substantially higher CTQ sum scores than non-maltreated participants (p < .001).

**Table 1:** Descriptive sample statistics stratified by attachment style and childhood maltreatment.

| Measure | Total | InsecCM- | InsecCM+<br>+ | SecCM- | SecCM+ | p |
| --- | --- | --- | --- | --- | --- | --- |
| **Demographic measures** |  |  |  |  |  |  |
| N (%) | 1317<br>(100) | 270<br>(20.5) | 300 (22.8) | 494<br>(37.5) | 253 (19.2) |  |
| Age (SD) | 34.93<br>(13.01) | 31.65<br>(11.10) | 35.45<br>(12.73) | 33.51<br>(12.98) | 40.58<br>(13.50) | <.001* |
| Sex female/male<br>(% female) | 867/450<br>(66%) | 169/101<br>(63%) | 202/98<br>(67%) | 319/175<br>(65%) | 177/76<br>(70%) | 0.300 |
| Monthly Household<br>Income in Euro<br>(SD) | 2,043.76<br>(1,730.38 ) | 1,938.00<br>(1,537.69 ) | 1,963.99<br>(1,619.10) | 2,388.28<br>(1,888.45 ) | 2,314.29<br>(1,532.57) | 0.007* |
| Years of Education | 13.85<br>(2.62) | 13.90<br>(2.58) | 13.38<br>(2.63) | 14.22<br>(2.53) | 13.59<br>(2.73) | <.001* |
| **Clinical Characteristics** |  |  |  |  |  |  |
| Diagnostic group<br>n HC/MDD | 896/421 | 191/79 | 106/194 | 437/57 | 162/92 | <.001* |
| MDD only,<br>Remission:<br>Partially/fully<br>remitted | 184/237 | 32/47 | 98/96 | 19/38 | 35/56 | 0.058 |
| MDD only,<br>Antidepressant:<br>No/Yes | 226/195 | 42/37 | 95/99 | 37/20 | 52/39 | 0.167 |
| MDD only,<br>Sackeim Index:<br>Medication Load | 1.38<br>(1.82) | 1.37<br>(1.88) | 1.52<br>(1.79) | 1.11<br>(1.93) | 1.24<br>(1.76) | 0.748 |
| **Attachment Measures** |  |  |  |  |  |  |
| RSQ Fear of<br>Abandonment | 2.57<br>(0.65) | 2.97<br>(0.63) | 3.02<br>(0.71) | 2.24<br>(0.37) | 2.26<br>(0.41) | <.001* |
| RSQ Fear of<br>Closeness | 2.25<br>(0.79) | 2.58<br>(0.76) | 2.93<br>(0.84) | 1.80<br>(0.47) | 1.98<br>(0.49) | <.001* |
| **Childhood Adversity** |  |  |  |  |  |  |
| CTQ Sum (SD) | 36.06<br>(11.93) | 30.16<br>(3.78) | 47.53<br>(13.08) | 28.54<br>(3.42) | 43.45<br>(11.94) | <.001* |
| **Neuroimaging Covariate** |  |  |  |  |  |  |
| Total Intracranial<br>Volume<br>(TIV) (SD) | 1,524.35<br>(147.57) | 1,533.64<br>(146.71) | 1,517.22<br>(160.54) | 1,537.69<br>(142.26) | 1,496.81<br>(138.96) | .002* |
| **Behavioral Outcomes** |  |  |  |  |  |  |
| Hamilton<br>Depression<br>Inventory (HAM-D) (SD) | 2.35<br>(3.08) | 2.63<br>(3.41) | 4.04<br>(3.74) | 1.15<br>(1.70) | 2.41<br>(2.92) | <.001* |
| Beck's<br>Depression<br>Inventory (BDI-I) (SD) | 6.52<br>(6.86) | 7.33<br>(7.02) | 11.66<br>(8.31) | 3.22<br>(3.75) | 6.03<br>(5.47) | <.001* |
| Perceived Stress<br>Scale (PSS) (SD) | 19.01<br>(8.80) | 20.92<br>(8.55) | 25.16<br>(8.74) | 14.48<br>(6.45) | 18.52<br>(8.03) | <.001* |
| Resilience Scale<br>(RS-25) (SD) | 134.53<br>(22.13) | 129.73<br>(22.02) | 119.12<br>(24.62) | 144.53<br>(15.71) | 138.41<br>(17.82) | <.001* |
| Global<br>Assessment of<br>Functioning (GAF)<br>(SD) | 85.41<br>(12.76) | 85.32<br>(12.36) | 76.79<br>(13.98) | 90.92<br>(8.70) | 85.07<br>(12.55) | <.001* |
| Global Symptom<br>Severity (SCL-90-R) (SD) | 0.38<br>(0.40) | 0.43<br>(0.39) | 0.70<br>(0.50) | 0.18<br>(0.18) | 0.34<br>(0.31) | <.001* |
| Social Support<br>(FSozU) (SD) | 4.36<br>(0.66) | 4.33<br>(0.57) | 3.83<br>(0.83) | 4.65<br>(0.40) | 4.43<br>(0.53) | <.001* |

**Note:** InsecCM-: Group with insecure attachment and no childhood maltreatment, InsecCM+: Group with insecure attachment style and history of childhood maltreatment, SecCM-: Group with secure attachment and no childhood maltreatment. SecCM+: Group with secure attachment and history of childhood maltreatment, "interpersonally resilient" group. Mean (*standard deviation*), TIV = total intracranial volume, HC = healthy control, MDD = major depressive disorder, CTQ = childhood trauma questionnaire, RSQ = Relationship scales questionnaire, HAM-D = Hamilton rating scale for depression, BDI = Beck depression inventory, PSS = perceived stress scale, RS-25 = resilience scale, GAF = global assessment of functioning, SCL-90-R = symptom checklist 90 revised.

### Main and interaction effects on behavioral outcomes

We conducted MANCOVA controlling for age, sex, and site to examine the main effects of childhood maltreatment (CM), attachment style, and their interaction on six behavioral outcomes: perceived stress (PSS), resilience (RS-25), global symptom severity (SCL-90), rater-based depressive symptoms (HAM-D), self-reported depressive symptoms (BDI), and global assessment of functioning (GAF).

### Significant main effect of childhood maltreatment on behavior

As expected, childhood maltreatment showed a significant multivariate effect on the combined behavioral outcomes: *F*(6, 1274) = 24.54, *p* < .001, Wilks’ Λ = .896, partial η2 = .104). Follow-up ANCOVAs revealed significant effects for all measures (all *p* < .001), indicating higher symptom severity and perceived stress, and lower resilience and global functioning among maltreated individuals (Table 2).

**Table 2:**
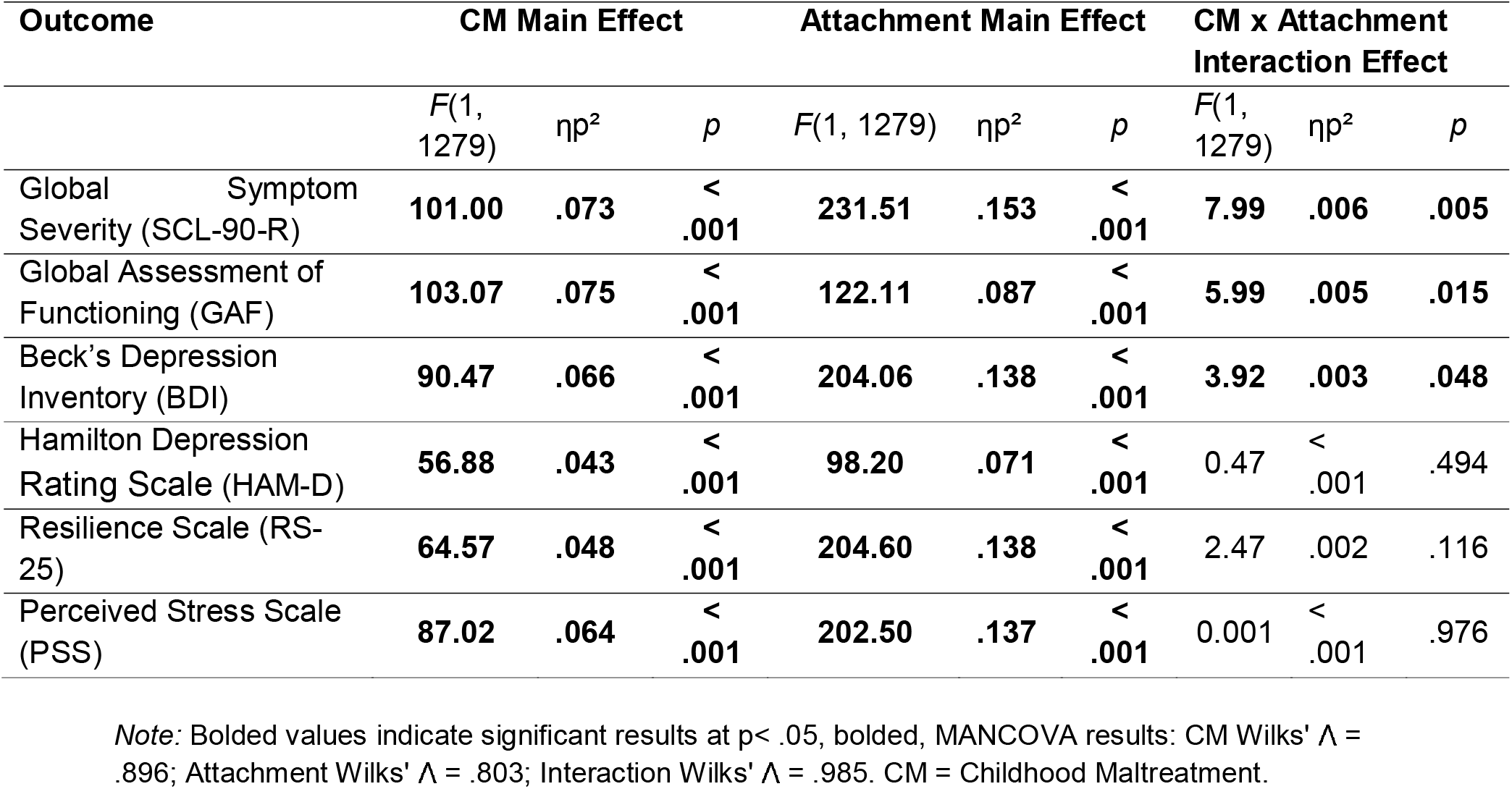
Follow-Up ANCOVA Results for Main and Interaction Effects on Behavioral Outcomes.

### Significant main effect of attachment on behavior

Attachment style also showed a significant multivariate effect (F(6,1274) = 52.16, p < .001, Wilks’ Λ = .803, partial η² = .197). Follow-up ANCOVAs revealed significant differences across all behavioral and clinical outcomes (all *p* < .001), with insecurely attached individuals reporting higher psychopathology and perceived stress as well as lower resilience and global functioning (Table 2).

### Significant interaction SCL-90-R, GAF, and BDI

A significant multivariate interaction of childhood maltreatment and attachment style was observed *F*(6,1274) = 3.21, *p* = .004, Wilks’ Λ = .985, partial η2 = .015. Post-hoc ANCOVAs revealed significant interaction effects on global symptom severity (SCL-90-R), global functioning (GAF) and self-reported depressive symptoms (BDI-I) (all *p*s < .05) but not for HAM-D, RS-25, or PSS (all *p*s < .05) (Table 2).

To further characterize these interaction effects, post-hoc pairwise comparisons of the estimated marginal means were conducted. Among individuals with childhood maltreatment, securely attached participants showed lower self-reported depressive symptoms compared to insecurely attached participants (BDI: SecCM+: *M* = 5.74, *SE* = 0.36 vs. InsecCM+: *M* = 11.5, *SE* = 0.35, *p* < .001). A similar pattern was observed for global symptom severity (SCL-90-R: SecCM+: *M* = 0.33, *SE* = 0.02 vs. InsecCM+: *M* = 0.69, *SE* = 0.02, *p* < .001) and global functioning (GAF: SecCM+: *M* = 85.31, *SE* = 0.74 vs. InsecCM+: *M* = 76.61, *SE* = 0.67, *p* < .001). These effects are illustrated in Figure 1.

**Figure 1.**
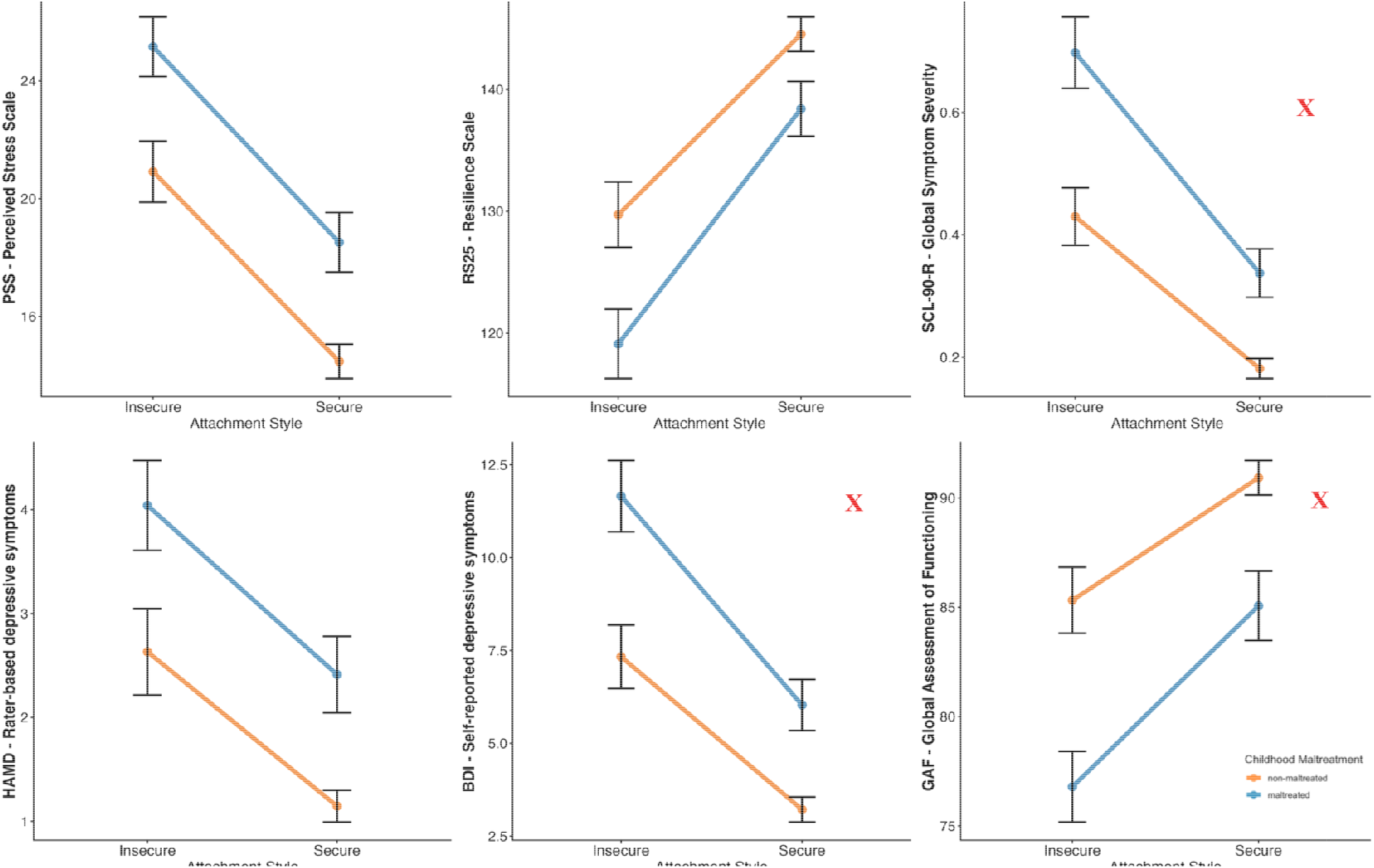
Main and interaction effects of attachment style and childhood maltreatment on all behavioral and clinical outcomes. *Note:* Mean scores for six behavioral and clinical measures across attachment style and childhood maltreatment groups. Error bars represent two standard errors of the mean. Significant childhood maltreatment × attachment interactions were observed for global symptom severity (SCL-90-R), self-reported depressive symptoms (BDI), and global functioning (GAF), indicated by red check marks.

In sum, both childhood maltreatment and attachment style were associated with significant differences across all behavioral and clinical outcomes. Further, observed interaction effects specifically affected self-reported depressive symptoms, global symptom severity, and global functioning, indicating more favorable outcomes among securely attached individuals with childhood maltreatment compared to insecurely attached individuals.

### Brain Structural findings reveal interaction effect of Childhood Maltreatment x Attachment

Whole-brain voxel-based morphometry analyses were conducted to examine the main effects of childhood maltreatment, attachment style, and their interaction on gray matter volume (GMV), while controlling for age, sex, TIV, site, and body coil change. Results were considered significant at cluster-level, corrected using FWE.

### No significant main effect of Childhood Maltreatment on brain structure

There was no significant effect for the positive or negative contrast of the main effect of childhood maltreatment on GMV.

### No significant main effect of Attachment on brain structure

There was no significant effect for the positive or negative contrast of the main effect of attachment style on GMV.

### Significant negative Interaction effect

A significant interaction between childhood maltreatment and attachment style was observed in the negative contrast (*T* = 4.55, *x/y/z* = - 64/-39/26, *k* = 951, *p* = .037, cluster-level FWE corrected), encompassing a cluster located in the supramarginal gyrus (Figure 2). No significant clusters were detected for the positive interaction contrast (*T* = 3.65, *x/y/z* = 36/42/40, *k* = 62, *p* = .951).

**Figure 2.**
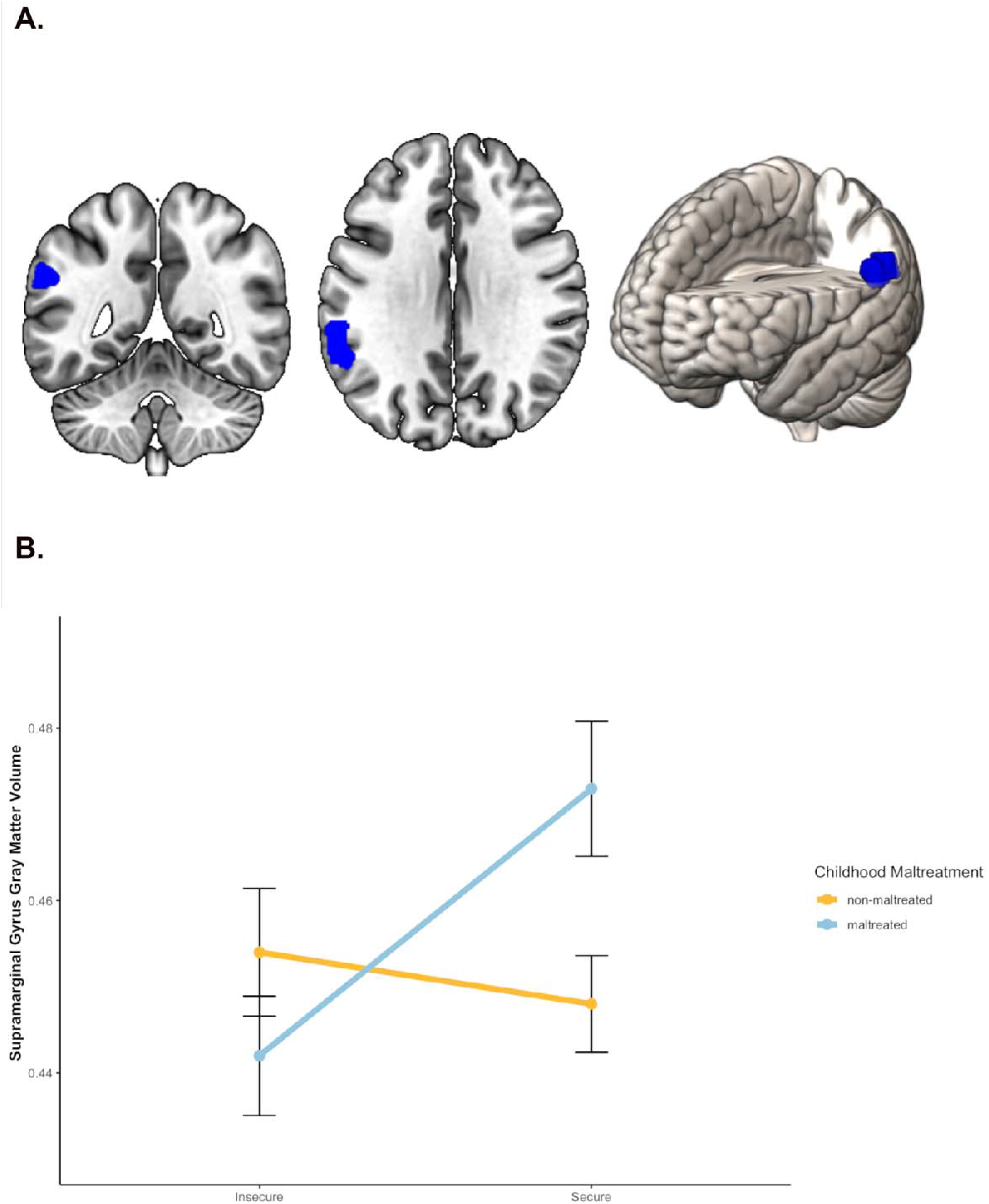
Significant interaction effect of childhood maltreatment x attachment in the left supramarginal gyrus. *Note*: A. Brain visualization of significant cluster, B. Attachment × Childhood Maltreatment interaction on gray matter volume by group, extracted eigenvariate cluster values depicted (values adjusted for covariates).

To further characterize the significant interaction, cluster eigenvariate values were extracted for each participant and analyzed using ANCOVA while controlling for the same covariates. These analyses indicated that individuals with childhood maltreatment and secure attachment showed larger GMV in the supramarginal gyrus cluster compared tp the other groups (Figure 2).

Additional models including medication load (Sackeim Index), antidepressant intake, and diagnostic group as covariates yielded comparable results, and the interaction effect remained statistically significant (see Supplement). A complementary dimensional moderation analysis using continuous childhood trauma severity (CTQ sum score) and attachment insecurity (multiplication term of RSQ subscales fear of closeness and fear of abandonment) revealed a significant interaction predicting cluster eigenvariate values, consistent with the categorical interaction observed in the main analysis (see Supplemental Analyses).

In sum, whole-brain analyses revealed a significant childhood maltreatment × attachment interaction on gray matter volume in the left supramarginal gyrus, with no significant main effects. The interaction remained significant in additional sensitivity analyses and was replicated in the dimensional moderation analysis.

## Discussion

In this study, we investigated whether secure attachment in the context of childhood maltreatment, conceptualized here as interpersonal resilience, is associated with differences in behavioral outcomes and brain structure. Using a large sample of 1,317 adults with and without a history of major depression, we observed main effects of childhood maltreatment and attachment style on multiple domains of mental health. Notably, we found evidence for a significant interaction between maltreatment history and attachment style on depressive symptoms, global functioning, and global symptom severity. These findings indicate that secure attachment may attenuate the negative consequences associated with childhood maltreatment.

On the neural level, this interpersonal resilience pattern was reflected in a significant interaction in the left supramarginal gyrus, where the interpersonally resilient group showed larger gray matter volume compared to all other groups. Importantly, this interaction remained significant in additional sensitivity analyses separately accounting for medication load, antidepressant intake, and diagnostic group, and was supported by a complementary dimensional analysis. Taken together, these findings support the conceptualization of secure attachment as a core component of interpersonal resilience following childhood maltreatment at both psychological and neuroanatomical levels.

As anticipated, we demonstrated significant main effects of childhood maltreatment and attachment on the behavioral outcomes. Here, securely attached individuals had better mental health outcomes compared to insecurely attached individuals; and non-maltreated individuals had better outcomes compared to maltreated individuals, including lower perceived stress, higher resilience, lower global symptom severity, lower self- and rater-based depressive symptoms. Our findings align with a large literature indicating that early interpersonal trauma confers broad vulnerability for elevated depressive symptoms, reduced resilience, higher perceived stress, and lower functioning ^8,45,46^.

These behavioral effects were evident across all phenotypic measures, even in a sample where acute depressive episodes were excluded, highlighting that the consequences of childhood maltreatment persist beyond periods of acute clinical illness. Likewise, the main effects of attachment style reinforce evidence that secure attachment is associated with fewer symptoms, higher resilience, and more adaptive stress regulation ^4,18^. Importantly, our findings showed larger effect sizes for attachment style compared to childhood maltreatment across all behavioral outcomes (ηp² = .071-.153 vs. .043-.075, respectively; Table 2), underscoring the importance of considering attachment as a relevant variable in mental health.

The detected behavioral interaction effects go beyond mere additive/subtractive effects and indicate that secure attachment modifies the association between childhood maltreatment and adult functioning and depressive symptom severity. Maltreated individuals with insecure attachment reported the highest symptom burden, whereas individuals with childhood maltreatment who maintained secure attachment showed significantly lower depressive symptoms, lower global symptom severity, and higher global functioning. Notably, this moderation pattern did not extend to all phenotypic domains, suggesting that attachment-related resilience may be particularly relevant for clinical symptom burden and overall functioning. Together, these results support the conceptualization of secure attachment despite childhood maltreatment as a form of interpersonal resilience.

In this context, interpersonally resilient individuals may retain a preserved or restored capacity for trusting relationships, adaptive help seeking, and effective emotion regulation - skills that have been repeatedly linked to lower vulnerability to depression and stress-related psychopathology ^47^. These skills likely also facilitate greater access to and engagement with social support, which may in turn reinforce secure attachment representations, suggesting a mutually reinforcing relationship between attachment security and social support rather than a strictly unidirectional pathway. This interpretation aligns with models suggesting that supportive relationships in adulthood may update internal working models, thereby buffering long-term effects of early adversity ^48^.

We identified a significant childhood maltreatment by attachment interaction in the left supramarginal gyrus. This region forms part of the inferior parietal lobule and is functionally heterogenous, contributing to phonological working memory, visual word recognition, and action understanding within the parietal component of the mirror neuron system ^49,50^. Beyond these functions, the supramarginal gyrus is consistently implicated in social cognitive processes, including cognitive empathy, perspective taking, and the ability to distinguish between one’s own and another person’s emotional state ^51–54^. Importantly, socio-affective and mentalizing-based training has been shown to produce structural plasticity in this region, including increases in cortical thickness (Valk et al., 2017), underscoring its relevance for interpersonal competencies. The finding that the interpersonally resilient group showed the largest gray matter volume in this area suggests that enhanced structural integrity in this social cognitive hub may support more adaptive functioning in close relationships after early adversity. Rather than exhibiting the volumetric reductions sometimes associated with maltreatment, this group displayed a pattern more consistent with a neural signature of adaptation.

We did not observe main effects of childhood maltreatment or attachment style on gray matter volume. While childhood maltreatment has been associated with widespread structural alterations, findings are inconsistent and moderated by multiple factors such as age, diagnosis and remission status, type and chronicity of adversity, and symptom severity ^20,56^. Recent work has demonstrated little overlap in childhood maltreatment findings across large partially overlapping cohorts ^57^. Similarly, the lack of a main effect of attachment may indicate that structural correlates of attachment variation vary and are heterogenous and difficult to detect independently of adversity context. Instead, our results suggest that neuroanatomical differences emerge most clearly when examining how attachment interacts with maltreatment to shape distinct pathways of vulnerability or resilience.

Several limitations should be considered. The assessment of childhood maltreatment relied on retrospective self-report, which may be affected by recall bias or mood-congruent memory ^58^. Both childhood maltreatment and attachment were assessed using self-report measures, which may introduce shared method variance. The attachment classification in this study relied on self-report cut-offs rather than interview-based assessments, which limits the ability to distinguish between specific insecure subtypes (i.e., anxious, avoidant, disorganized). Grouping them together therefore captures the central distinction between secure and insecure attachment relevant to the mechanisms examined here, even if some subtype-specific nuances are not represented ^25,59,60^. Additionally, we did not account for occurences of post-traumatic stress disorder (PTSD), which may have influenced the behavioral and neural differences observed here ^61,62^. Further, VBM findings can be affected by misalignment during normalization, tissue misclassification, and individual variability in cortical folding and thickness ^63^. Lastly, the cross-sectional design of the study limits causal inference regarding the temporal relationship between childhood maltreatment, attachment style, behavioral symptoms and brain structure.

Our findings have several theoretical and clinical implications. First, they support the value of moving beyond risk-focused frameworks toward models that identify protective processes. Even among individuals exposed to significant adversity, secure attachment appears to confer psychological and neurobiological advantages. Second, the results underline the relevance of attachment-based interventions. Therapeutic approaches that strengthen secure relational bonds, enhance trust, and improve mentalizing may help shift trajectories after adversity. If attachment security corresponds to measurable neural differences in regions central for social cognition, interventions targeting these capacities may support adaptive changes in social cognitive functioning. Third, the present findings reinforce the importance of dimensional approaches that include individuals across diagnostic boundaries. Both attachment style and depressive symptoms varied continuously across the sample, and resilience processes were detectable beyond categorical diagnoses.

In sum, we demonstrate that secure attachment may buffer the psychological and neural consequences of childhood maltreatment. Across individuals with and without a history of depression, we showed that secure attachment is associated with fewer self-reported and rater-based depressive symptoms, higher psychometric resilience and global functioning, and lower symptom severity and perceived stress, with comparatively larger effect sizes than those for childhood maltreatment. Further, adults who experienced maltreatment but maintain secure attachment show larger gray matter volume in the left supramarginal gyrus, a region central for social cognition and self-other distinction. These results highlight secure attachment as a protective factor and suggest potential pathways through which supportive relationships may support interpersonal resilience and adaptive development after early adversity. Incorporating protective factors into research is essential for understanding why individuals with similar histories of adversity show different outcomes and for identifying mechanisms that may inform prevention and intervention efforts.

## Acknowledgments

We are deeply indebted to all study participants and staff. A list of further acknowledgments can be found at: for2107.de/acknowledgements.

During the preparation of this work the authors used LLM to improve readability and language. After using this tool, the authors reviewed and edited the content as needed and take full responsibility for the content of the publication.

