## Supplemental Material for "Secure Attachment despite Childhood Maltreatment: Behavioral and Neural Correlates of Interpersonal Resilience"

Brosch *et al.*

**Correlation Matrix of Behavioral and Clinical Measures**

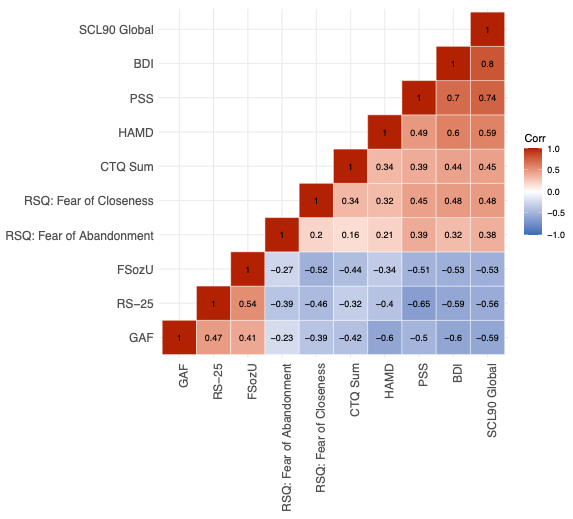

**Supplementary Figure 1: Behavioral and clinical measures are moderately to highly correlated.**

The 2D grids display pairwise Pearson correlation coefficients between behavioral measures. SCL90 Global = SCL-90-R Global Severity Index, BDI = Beck’s Depression Inventory, PSS = Perceived Stress Scale, HAMD = Hamilton Depression Rating Scale, CTQ Sum = Childhood Maltreatment Sum Score, RSQ = Relationship Scales Questionnaire, FSozU = Perceived Social Support, RS-25 = Self-reported Resilience, GAF = Global Assessment of Functioning.

**Behavioral Scores by group**

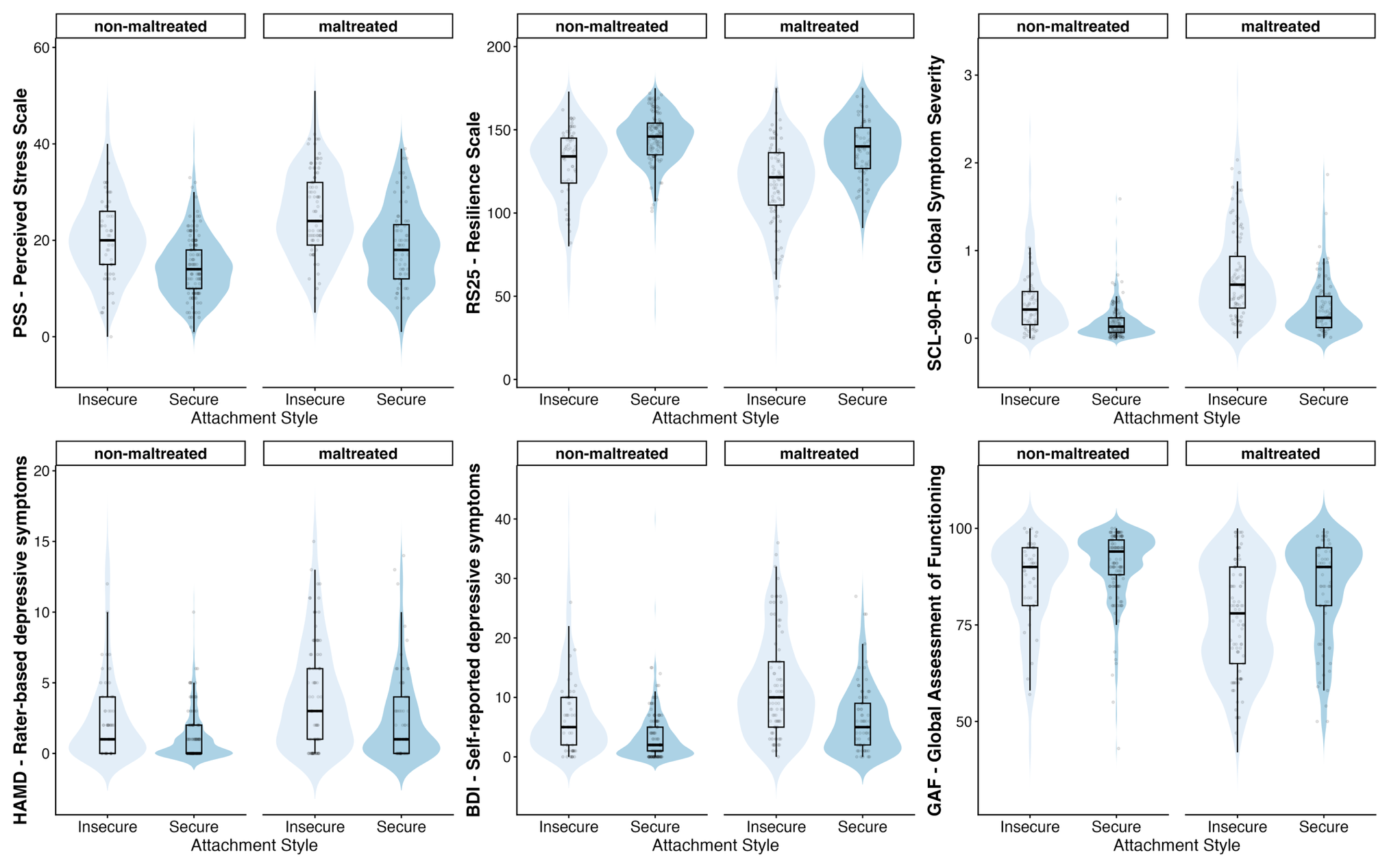

**Supplementary Figure 2. Violin-Boxplot Representation of Behavioral Outcomes Across Groups.**

Violin plots display the full distribution of individual scores within each attachment style and maltreatment group. Embedded boxplots represent the median and interquartile range. To reduce overplotting, a random 25% subsample of individual observations is shown as jittered points.

**Supplemental Analyses**

To assess whether the interaction effect identified in the voxel-wise analysis was robust to potential confounders, we extracted subject-level eigenvariate values from the significant cluster and tested whether the **childhood maltreatment × attachment** interaction remained significant when additionally covarying for dichotomous antidepressant intake, medication load (Sackeim Index), or group. Medication analyses were restricted to the MDD group, as antidepressant intake and medication load were not applicable to healthy controls.

**Dichotomous Antidepressant Intake**

In the MDD sample, the interaction between childhood maltreatment and attachment style remained significant after additionally adjusting for antidepressant intake, age, sex, site, and total intracranial volume (TIV), (β = 0.048, SE = 0.011, t(412) = 4.28, p < .001). Antidepressant intake itself was not significantly associated with the outcome (β = -0.004, p = .455). The overall model was significant, F(8, 412) = 55.53, p < .001.

**Sackeim Index Medication Load**

**Similarly, the interaction between childhood maltreatment and attachment style remained significant after additionally adjusting for medication load (Sackeim Index), age, sex, site, and TIV in the MDD group** (β = 0.048, SE = 0.011, t(412) = 4.28, p < .001). The Sackeim Index was not significantly associated with the outcome (β = 0.00014, p = .396). The overall model was significant, (F(8, 412) = 55.58, p < .001).

Together, these findings suggest that the interaction effect is robust to adjustment for antidepressant intake and medication load.

**Group**

To test whether group (HC/MDD) would impact our brain imaging findings, we added the group variable as a covariate. In the full sample, the interaction between childhood maltreatment and attachment style remained significant after adjusting for diagnostic group, age, sex, site, and TIV (β = 0.037, SE = 0.006, t(1308) = 6.35, p < .001). Diagnostic group was also independently associated with the outcome (β = -0.007, p = .036). The overall model was significant, F(8, 1308) = 181.10, p < .001.

This indicates that diagnostic group is associated with overall differences in the outcome, while the detected interaction effect of childhood maltreatment and attachment style on brain remains statistically significant after accounting for group.

**Dimensional Analysis**

To complement the categorical CM × attachment analysis, we conducted a secondary dimensional moderation analysis using subject-level eigenvariate values extracted from the significant cluster. A multiple linear regression model tested whether continuous childhood trauma severity (CTQ sum score) interacted interacted with attachment insecurity **(RSQ composite score of fear of closeness and fear of abandonment)** to predict eigenvariate cluster values, controlling for age, sex, site, and total intracranial volume (TIV).

In the dimensional model, the interaction between childhood maltreatment severity (CTQ sum score) and attachment insecurity was significant, β = −<.001, SE = <.001, t(1309) = −2.06, p = .040. The overall model was significant, F(7, 1309) = 195.4, p < .001.

**These results are consistent with the categorical interaction analysis and indicate that attachment insecurity moderates the association between childhood trauma severity and gray matter volume in the identified cluster.**
